# The role of CYP3A-CYP2E1 interactions in activation of CYP3A enzymes by chronic alcohol exposure

**DOI:** 10.64898/2026.02.06.703602

**Authors:** Dmitri R. Davydov, Kannapiran Ponraj, Nadezhda Davydova, Guihua Yue, Dilip Kumar Singh, Arpita Guha Neogi, Kari A. Gaither, Bhagwat Prasad

**Affiliations:** Department of Chemistry, Washington State University, Pullman, WA, 99164; Division of Translational and Clinical Pharmacology, Cincinnati Children’s Hospital Medical Center, Cincinnati, OH, 45229; Department of Pharmaceutical Sciences, Washington State University, Spokane, WA, 99202

**Keywords:** cytochrome P450, alcohol, drug metabolism, protein-protein interaction, protein cross-linking, CYP2E1, CYP3A4, 7-benzyloxyquinoline, ivermectin, human liver microsomes, CYP3A5, NADPH-cytochrome P450 reductase, protein-protein interactions, alcohol consumption, alcohol-drug interactions, crosslinking mass spectrometry

## Abstract

Aiming to examine the effect of chronic alcohol exposure on the activity of CYP3A enzymes in human liver, we studied the metabolism of two CYP3A-specific substrates, 7-benzyloxyquinoline (7-BQ) and ivermectin, in 23 preparations of human liver microsomes (HLM) obtained from donors with documented alcohol exposure, graded from non-drinkers to heavy alcoholics. All HLM samples were characterized for the composition of the cytochrome P450 pool and the abundances of other drug-metabolizing and endoplasmic reticulum-stress-related enzymes by global proteomics. Our studies revealed a striking increase in the activities of CYP3A enzymes caused by chronic alcohol exposure. This effect is not associated with CYP3A enzyme levels, which do not correlate with alcohol exposure. Instead, the rates of 7-BQ and ivermectin metabolism correlate with the content of alcohol-inducible CYP2E1. However, this enzyme does not metabolize ivermectin, and its activity with 7-BQ is negligible. These results suggest that the observed acceleration of the elimination of drugs metabolized by CYP3A enzymes by alcohol exposure is due to functional effects of the interaction between CYP3A and CYP2E1. To elucidate the potential mechanism of this effect, we studied the formation of CYP2E1-CYP3A4 complexes in CYP3A4-containing Supersomes with co-incorporated CYP2E1 using tag-transfer chemical crosslinking mass spectrometry (CX-MS). These experiments confirmed physical interactions between the proteins and allowed the identification of CYP3A4 residues at the sites of contact. This information was used to build structural models of the CYP2E1-CYP3A4 complex and to propose possible mechanisms for the observed effects.

## 1. Introduction

An essential part of alcohol-related fatalities is associated with deleterious alcohol interactions with drugs, including both prescribed medications and drugs of abuse. The list of drugs whose therapeutic properties are changed by alcohol consumption comprises a variety of commonly prescribed medications [1, 2, 3, 4]. In most cases, chronic alcohol exposure increases drug clearance [3, 5, 6, 7].

Understanding the mechanisms of alcohol-drug interactions is an ultimate prerequisite for developing optimal practices for medical treatment in alcohol consumers. It is essential for the prevention of deleterious effects of alcohol-drug interactions (ADI) and the identification of potentially important but yet unrecognized effects of alcohol on drug properties. A critical role in ADI is played by the human drug-metabolizing cytochromes P450. Alcohol exposure induces distinct and isoform-specific changes in P450 expression. The multifold increase in cytochrome P450 2E1 (CYP2E1) content in the liver and other tissues is one of the most important effects of alcohol on protein expression [8, 9, 10]. Nonetheless, the involvement of CYP2E1 in pharmacokinetic interactions of alcohol is commonly considered insignificant due to the presumed minimal role of CYP2E1 in drug metabolism [4, 5]. Our recent proteomic study on a large collection of human liver microsomes (HLM) from donors with various degrees of alcohol exposure, besides confirming alcohol-induced expression of CYP2E1, also revealed upregulation of NADPH-cytochrome P450 reductase (CPR), CYP2B6, and CYP2J2, as well as downregulation of CYP1A2, CYP2C8, CYP2C9, CYP4A11, and cytochrome *b*_5_ [11].

Besides acetaminophen, which is metabolized by alcohol-inducible CYP2E1 [12], pronounced effects of alcohol exposure on pharmacokinetics were observed with anti-depressants imipramine, the substrate of CYP2C19 [13, 14], and desipramine metabolized by CYP2D6 [14], or anticoagulant warfarin metabolized by CYP1A2 and CYP2C9 [6, 15]. Among the drugs involved in ADI, the longest is the list of CYP3A substrates. It includes antibiotic doxycycline [16], sedative agents diazepam [3, 17] and chlordiazepoxide [18, 19], barbiturate thiopentone (3A4) [20], opiates methadone [21, 22] and fentanyl [23], antidepressant amitriptyline [24, 25], and anticonvulsants carbamazepine [26] and phenytoin [6, 27].

Despite earlier reports evidencing an ethanol-induced increase in CYP3A4 expression in cell culture and animal models [28, 29], we did not observe a correlation between the level of alcohol consumption and the abundance of CYP3A enzymes in the human liver [11]. According to our results, chronic alcohol exposure does not affect the expression level of CYP3A4. This finding seemingly contradicts numerous indications of the effect of alcohol exposure on CYP3A-dependent drug metabolism cited above.

However, the impact of alcohol on drug metabolism may stretch far beyond the drugs metabolized by P450 species affected by alcohol consumption. According to our concept, the ensemble of P450 enzymes in the endoplasmic reticulum of liver cells functions as an integrated multienzyme system, in which its members compete for a common electron donor (CPR) and interact to form heteromeric complexes, thereby affecting their function [30]. Consequently, alcohol-induced changes in P450 expression alter the protein-protein interaction landscape and induce a wide range of functional changes [30, 31].

To probe this hypothesis in the present study, we investigated the metabolism of two CYP3A-specific substrates, the fluorogenic substrate 7-benzyloxyquinoline (7-BQ) and the antiparasitic drug ivermectin, using a collection of 23 proteomically characterized HLM preparations from donors with varying degrees of alcohol exposure. Our studies demonstrated a strict correlation between CYP3A activity and the abundance of alcohol-inducible CYP2E1 protein in the human liver. These results confirm our previous studies, in which we revealed an increase in the rate of 7-BQ metabolism in pooled HLM preparations after enrichment with CYP2E1 by incorporating the purified enzyme into their membranes [32].

To explore the molecular mechanisms underlying the observed effects, we studied the interactions between CYP2E1 and CYP3A4 using tag-transfer chemical crosslinking mass spectrometry (tag-transfer CX-MS). These studies confirmed the formation of mixed oligomers of the two enzymes and provided information for modeling the architecture of their complex. Collectively, our results demonstrate an increase in CYP3A activity caused by chronic alcohol exposure, which is mediated by functional effects of CYP3A-CYP2E1 interactions.

## 2. Materials and Methods

### 2.1. Chemicals

7-Benzyloxyquinoline (7-BQ) was purchased from Toronto Research Chemicals (Vaughan, ON, Canada). Before use, it was further purified to remove traces of contaminating 7-hydroxyquinoline (7-HQ) by silica chromatography as described in section 2.6. 7-HQ was obtained from Acros Organics, a part of Thermo Fisher Scientific (New Jersey, NJ). Ivermectin was obtained from ApexBio (Boston, MA). Acetoacetanilide was procured from the Tokyo Chemical Industry (Tokyo, Japan). Glucose-6-phosphate dehydrogenase from Leuconostoc mesenteroides was obtained from Worthington Biochemical Corporation (Lakewood, NJ). NADP and Glucose-6-phosphate were the products of Millipore Sigma (Burlington, MA, USA). MS Grade Pierce™ Trypsin Protease, LC-MS grade acetonitrile, formic acid, dithiothreitol (DTT), and iodoacetamide (IAA) were purchased from Fisher Scientific (Fair Lawn, NJ). All other reagents were of ACS grade and used without additional purification.

### 2.2. Protein expression and purification

N-terminally truncated (Δ3-20) and C-terminally His-tagged CYP2E1 [33] was expressed in *E. coli* TOPP3 cells and purified as described earlier [34].

### 2.3. HLM samples from individual donors

In this study, we used 23 microsomal preparations from liver specimens obtained from non-identifiable individual donors. The set comprises 17 individual liver samples from the biobank established in the Prasad Laboratory and six samples from donors with a history of moderate-to-heavy alcohol consumption procured from BioIVT Corporation (Baltimore, MD). The liver samples were included based on documented alcohol intake history. The demographic characteristics of the donors are described in our previous publication [35]. Preparation of the HLM fraction was performed as described earlier [35].

All HLM preparations were subjected to the determination of relative abundances of the individual cytochrome P450 species, their redox partners (CPR and cytochrome *b*_5_), and protein markers of alcohol exposure (HSPA5, PDIA3, P4HB, and CES2 [11]). This analysis was performed using the global proteomics-based Total Protein Approach (TPA) as described previously [35, 36]. Its results are available in the dataset on Mendeley Data [37].

### 2.4. Microsomes containing recombinant human cytochromes P450

Most of the preparations of insect cell microsomes containing baculovirus-expressed individual P450 enzymes (Supersomes_TM_) and human CPR were produced by Gentest (Huntsville, AL), now a part of Discovery Life Sciences (Woburn, MA). In the present study, we used the preparations containing CYP2B6 (lot 31487), CYP2C8 (lot 81760), CYP2C9 (lot 75854), CYP2C19 (lot 73445), CYP2E1 (lot 23012), CYP3A4 (lot 81745), CYP3A5 (lot 89573), and CYP4A11 (lot 2402280). All those preparations except for the CYP1A2 and CYP4A11 Supersomes also contained human cytochrome *b*_5_ co-expressed. The preparations of insect cell microsomes containing human CYP2A6 (lot 2520331) and CYP2D6 (lot 2350106), along with human CPR and cytochrome *b*_5_ (Baculosomes®), were procured from Thermo Fisher Scientific (Waltham, MA, USA).

### 2.5. Characterization of the content of protein, NADPH-cytochrome P450 reductase, and cytochromes P450 and b_5_ *in HLM preparations*

Protein concentrations in microsomal suspensions were determined by the bicinchoninic acid assay. The concentration of CPR in microsomal membranes was determined based on the rate of NADPH-dependent reduction of cytochrome *c* at 25 °C, and the effective molar concentration of CPR was estimated using the turnover number of 3750 min^-1^ [34]. The total concentration of cytochromes P450 in HLM was determined with a variant of the “oxidized CO versus reduced CO difference spectrum” method described earlier [34].

### 2.6. High-throughput fluorimetric assays of 7-BQ and ivermectin metabolism

The process of developing the 7-BQ procedure is outlined in section 3.1 under Results. As indicated there, a critical difficulty for microplate-based acquisition of substrate saturation profiles of 7-BQ has been contamination of commercially available 7-BQ preparations with traces of the product of its debenzylation, 7-HQ. To address this issue, we purified 7-BQ by chromatography on a Normal Phase Silica Solid Phase Extraction Column. To this aim, 5 mg of 7-BQ dissolved in 0.5 ml chloroform were passed through a BJ9050 100 mg Silica column (Honeywell Burdick & Jackson, Muskegon, MI, USA). The column retained all the contaminating 7-HQ. The column was then washed with 1 ml of chloroform. The flow-through liquid, combined with the wash liquid, was then subjected to chloroform evaporation under a stream of argon gas. The purified 7-BQ was dissolved in 0.5 ml acetonitrile, and the concentration of the resulting stock solution was determined from its absorbance at 324 nm using the extinction coefficient of 4.43 mM^-1^cm^-1^.

The acquisition of the substrate saturation profiles (SSPs) for 7-BQ and ivermectin metabolism was performed using an automated microplate-based procedure with an OT-2 liquid-handling robot (Opentrons Inc., Brooklyn, NY) and a Cary Eclipse fluorometer equipped with a plate reader accessory (Agilent Technologies, Santa Clara, CA, USA). The assays with ivermectin were based on the fluorometric determination of formaldehyde (FA) formed in its O-demethylation and were carried out as described earlier [38]. The protocol for the assays with 7-BQ was similar to that described earlier for the studies with 3-(6-Methoxynaphthalen-2-yl)acrylic acid (MONACRA) [39] except that 2.3 M Na-glycine buffer, pH 10.4, was replaced with the same buffer at 0.4 M concentration. This solution was added at the end of the assay procedure to neutralize the trichloroacetic acid quenching reagent and bring the pH to 7–7.5, which is optimal for the fluorometric determination of 7-HQ.

All activity assays were performed in 0.1 M Na-HEPES buffer containing 60 mM KCl. Multi-well plates for these assays were prepared using 12 stock solutions of 7-BQ or ivermectin, obtained by serial dilution with a dilution factor of 1.73333 (≈√3). The stock solutions with the highest substrate concentration (2.31 and 0.4 mM for 7-BQ and ivermectin, respectively) were prepared in incubation buffer containing 10% acetonitrile. Further dilution was performed with the acetonitrile-free buffer. The maximum substrate concentration used in the series was 577 µM for 7-BQ and 100 µM for ivermectin. The incubation buffer also contained 100 µM NADP, 2 mM glucose-6-phosphate, and 1 unit/ml glucose-6-phosphate dehydrogenase. Incubation was performed in the heater-shaker module of the OT-2 robot at 29.5–31 °C and 1200 rpm shaking speed. The incubation times were 12 min and 30 min for the assays with 7-BQ and ivermectin, respectively.

After the end of the OT-2 run, the incubation plates were centrifuged at 3,600 rpm at room temperature in a Beckman Allegra 6R centrifuge with a GH-3.8 swing-out rotor with multiwell-plate adapters. The centrifugation time was 15 and 30 min for the 7-BQ and ivermectin assays, respectively. In the latter case, additional time was required to ensure completion of the Hantzsch reaction, which is used for the determination of formaldehyde.

For ivermectin metabolism assays, excitation spectra were acquired as described earlier for FA-based assays [38]. The assays for 7-HQ formation were based on the spectra of fluorescence with synchronous scanning of excitation and emission in the wavelength range of 290 – 460 nm with Δλ = 110 nm. The series of these spectra corresponding to 12 concentrations of 7-BQ was subjected to principal component analysis, and the amount of formed 7-HQ was determined based on the approximation of the first three principal components by the set of prototypical spectra of fluorescence of 7-HQ, 7-BQ, and NADPH.

In the assays with HLMs, the rate of enzyme turnover was calculated relative to the concentration of CPR, the limiting component in the human microsomal monooxygenase system. In the case of insect cell microsomes with recombinant P450 enzymes (Supersomes™ and Baculosomes®), turnover numbers were calculated relative to the P450 enzyme concentration, which is lower than that of CPR in these systems.

### 2.7. Global kinetic analysis of substrate saturation profiles

The datasets for global kinetic analysis were assembled as sets of averages of 4–7 SSP obtained with each HLM preparation. The assembled datasets were subjected to Principal Component Analysis (PCA). The PCA results were used to find a set of two Hill or Michaelis-Menten equations whose linear combinations best approximate each SSP trace. To find the parameters of these components, we used a Nelder-Mead optimization procedure combined with the SURFIT algorithm for multidimensional linear regression [40], applied to the first two principal component vectors and the average of all traces in the dataset (the base vector of the analysis). The optimized set of two kinetic components resolved thereby was used to determine the amplitudes of the two individual phases in each SSP using the Iterative Target Transform Factor Analysis (ITTFA) technique [41, 42]. Description of our algorithm designed for this study is available in the Zenodo repository (https://doi.org/10.5281/zenodo.19102227). The square correlation coefficients for the resulting approximations of the SSPs were always >0.98 and, in most cases, >0.995. The amplitudes of the individual phases of SSPs were then analyzed to probe their correlations with the composition of the cytochrome P450 ensemble in HLM samples.

All manipulations with the dataset, Principal Component Analysis, and regression analysis were performed using our SpectraLab software [43, 44], which is available for download at https://cyp3a4.chem.wsu.edu/spectralab.html.

### 2.8. Analysis of correlations between the SSPs and the composition of the P450 pool

To find the correlation of the shape and amplitudes of SSP with variations of the abundances of individual P450 enzymes, we calculated the vectors of relative abundance (VRA) for 15 major P450s (CYP1A2, CYP2A6, CYP2B6, CYP2C8, CYP2C9, CYP2C18, CYP2C19, CYP2D6, CYP2E1, CYP3A4, CYP3A5, CYP4A11, CYP4F2, CYP4F12, and CYP2J2) and supplemented them with the vectors of relative abundance of CPR and cytochrome *b*_5_. The procedure for these calculations is described in our recent publication [35], and the resulting set of VRA vectors for all 23 HLM samples used in this study can be found in the dataset available at Mendeley Data [37]

Analysis of correlations between the composition of the P450 ensemble and the shape and amplitude of SSP was based on approximating the profiles of the amplitudes of each kinetic component (*V*_max1_ and *V*_max2_) and their total (*V*_max total_) with linear combinations of the vectors of relative fractional abundance of eleven major cytochrome P450 species complemented with the vectors of relative abundance of CPR and cytochrome *b*_5_. For these trials, the vectors of the *V*_max_ values for the HLM samples in the dataset were fitted to linear combinations of one to four vectors of relative abundance using the SURFIT algorithm of multidimensional linear regression [40]. We wrote a script for the SpectraLab software that successively applies the SURFIT algorithm to all possible combinations of up to 4 vectors of relative abundances for the P450 species under analysis and identifies the best approximation (in terms of the squared correlation coefficient).

### 2.9. Probing CYP2E1-CYP3A4 interactions in microsomal membranes with tag-transfer chemical crosslinking mass spectrometry

#### 2.9.1 Design of CXMS experiments

In the design of CXMS experiments, we combined tag-transfer CXMS with the molecular fishing strategy. Cleavable photo-activated crosslinking agent used in our de-sign is trifluoromethyl phenyl diazirine methanethiosulfonate (TFMD-MTS, 3-{4-[(methanesulfonylsulfanyl)methyl]phenyl}-3-(trifluoromethyl)-3H-diazirine) [45]. Crosslinker-activated molecular bait was His-tagged N-terminal truncated CYP2E1 protein modified with TFMD-MTS in the molecular ratio 1:3. Incorporation of the protein into insect cell microsomes (Supersomes®) containing recombinant CYP3A4, CPR, and cytochrome *b*_5_ was followed by light exposure, solubilization of the membranes, and bait isolation by Ni-affinity chromatography. The isolated bait protein was subjected to SDS-PAGE, and the gel fragments corresponding to molecular masses 45-65 kDa (zone A), 65–120 kDa (zone B), and 120–300 kDa (zone C) were digested using an optimized protocol [36] and analyzed by nanoLC-HRMS.

#### 2.9.2 Modification of CYP2E1 with TFMD-MTS and its incorporation into CYP3A4-containing microsomes

TFMD-MTS labeling was performed in 0.1 M HEPES buffer, pH 7.4, containing 0.15 M KCl and 20% glycerol. Before modification, DTT present in the stock solutions of purified CYP2E1 was removed by passing the solutions through a Bio-Gel P6 (Bio-Rad, Hercules, CA, USA) spin-out column equilibrated with the reaction buffer. After diluting the sample to 10 µM, it was supplemented with 0.2% Igepal CO-630 (10% solution in the reaction buffer) and TFMD-MTS (20 mM in acetone) at a 3:1 molar ratio to the P450 protein. After saturating the sample with argon gas by gentle bubbling, it was incubated for 1 hour at room temperature under continuous stirring. After removal of the detergent by treatment with BioBeads SM2 resin (BioRad, Hercules, CA) and concentrating the protein to ~50 µM with an Amicon Pro 30kDa cutoff centrifugal concentrator (Millipore, MA), the sample was passed through a spin-out Bio-Gel P6 column equilibrated with the reaction buffer to remove unreacted label.

Suspension of CYP3A4-containing Supersomes™ supplied by the manufacturer (1µM CYP3A4) was centrifuged at 150,000 × g in an Optima TLX ultracentrifuge (Beckman Coulter Inc., Brea, CA, USA) with a TLA100.3 rotor for 90 min at 4°C and resuspended in 0.1 M Na-HEPES buffer containing 150 mM KCl and 0.25 M Sucrose (Microsomes Storage Buffer) to P450 concentration of 10 µM. The suspension was supplemented with TFMD-MTS-labeled CYP2E1 at a 1:1 molar ratio to the microsomal CYP3A4 protein. After saturating with argon gas, the mixture was incubated for 16–20 h at 4°C with continuous stirring. The suspension was then centrifuged at 150,000 × g as described above. The pellet was resuspended in the Microsome Storage Buffer to a P450 concentration of 10-15 µM.

#### 2.9.3. Photo-activated crosslinking, subsequent isolation of the bait protein, and its fractionation with SDS-PAGE

The suspension of HLM with incorporated TFMD-MTS-modified CYP2E1 was diluted to a P450 concentration of 5 µM by argon-saturated Microsome Storage Buffer and placed into a 1 × 1 cm optical quartz cell. The cell was flushed with argon gas, tightly sealed, and exposed to 375 nm light from the M375L4 LED light source (Thor Labs, Newton, NJ) at continuous stirring at 4 °C in a setup similar to that described by Horne et al. [45]. After one hour of light exposure, the suspension was centrifuged at 105,000× g for 90 min. The pellet was resuspended in 1 mL of 0.125 M K-phosphate buffer, pH 7.4, containing 20% glycerol and 0.5% Igepal CO-630. The mixture was incubated for 2 h at 4 °C under continuous stirring, then centrifuged at 105,000× g for 90 min. The supernatant was applied to 0.2 mL of HisPur™ Ni-NTA resin (Thermo Fisher Scientific, Waltham, MA, USA) placed into a small spin-out and equilibrated with the same buffer. After one hour of incubation of the closed column under periodic shaking, the column was washed with multiple subsequent 1 mL portions of the same buffer containing 0.5% CHAPS until the optical density of the flow-through at 280 nm decreased below 0.025. The bound protein was eluted with 500 mM K-Phosphate buffer, pH 7.4, containing 20% glycerol, 0.5% CHAPS, and 250 mM imidazole. The detergent was removed using a Bio-Beads SM-2 resin (Bio-Rad, Hercules, CA, USA). The protein solution was concentrated to 10–20 mg/mL using a Centrisart I MWCO 100 kDa concentrator (Sartorius AG, Göttingen, Germany).

The proteins extracted from Ni-NTA resin were subjected to SDS-PAGE on 4–15% Mini-PROTEAN® TGX™ Precast Protein Gels (Bio-Rad, Hercules, CA, USA). The Broad Multi-Color Pre-Stained Protein Standard from GenScript (Piscataway, NJ, USA) was used for calibration. The gels were stained with Coomassie Brilliant Blue R-250 Staining Solution (Bio-Rad) and subjected to fragmentation, as described above. The resulting gel fragments were subjected to trypsin digestion and then analyzed by global proteomics.

#### 2.9.4. Mass-spectroscopic analysis and identification of the TFMD-tagged peptides

Tryptic digest (1 µg) of each gel fragment was analyzed on the EASY-nLC 1200 series coupled with a Thermo Scientific Q-Exactive HF mass spectrometer (Thermo Fisher Scientific, Waltham, MA). Desalting of the digest containing both unlinked and cross-linked peptides was achieved using an Acclaim PepMap trap column (75 µm × 2 cm, 3 µm particle size) (Thermo Scientific, Part Number 164535) connected with the analytical PepMap RSLC C18 column (75 × 25 cm, 2 µm particle size) (Thermo Scientific, Part Number ES902), respectively. The mobile phase consisting of 0.1% formic acid in water (A) and 0.1% formic acid in 80% acetonitrile (B) was run in gradient mode (%B): 2–6% (0–5 min), 6–30% (5–60 min), 30–100% (60–65 min), and 100% (65–80 min) with a flow rate of 300 nL/min. The injection volume and column temperature were set to 1 µL and 40 °C, respectively. The MS data were acquired in positive-ion polarity using full-scan and data-dependent acquisition (DDA) modes. The scan range in full MS and DDA modes was kept m/z 350 to 1600 and m/z 200 to 2000, respectively. At m/z 200, the resolution for full MS and DDA was 120,000 and 30,000, respectively. Crosslinked peptides were analyzed using the Kojak version 3.4 crosslink identification platform [46] available at https://kojak-ms.systemsbiology.net/. Searches for crosslinked peptides were performed with optimized crosslinking parameters. The analysis aimed to investigate CYP3A4-tagged peptides created by the photoactivatable MTS-TFMD cross-linker. This crosslinking reaction produces crosslinked peptides with a distinct 261.0435 Da mass addition linked to the CYP3A4 tryptic peptides. Next, Kojak search results were processed with Percolator [47] to produce a statistically validated set of crosslinked peptide identifications at a 5% false discovery rate (FDR). Further, MS/MS fragmentation analysis of crosslinked peptides was visualized and confirmed by Proxl [48].

### 2.10. Protein docking

The docking procedure used to model the architecture of the CYP2E1-CYP3A4 complex, based on CXMS results, combined manual selection of a starting pose with subsequent restrained docking using HEX 8.0.0 interactive docking software [40], followed by restrained docking using the HADDOCK 2.4 web server [49, 50]. The procedure began with manual selection of the initial pose to ensure proper orientation of both molecules for membrane embedding and to position some of the surface-exposed cysteines of CYP2E1 (C261, C268, C480, and C488) in proximity to the TFMD-tagged regions of CYP3A4. This pose served as a starting point for restrained docking with HEX 8.0.0 [51, 52]. As restraints, we used a 15° turning-angle range for both the receptor (CYP2E1) and the ligand (CYP3A4), and a 0° twist range. Docking was based on the combined optimization of shape and DARS potential [49]. The best pose compatible with membrane incorporation of both proteins was selected as the starting point for restrained docking with HADDOCK 2.4. As active residues, we specified the CYP2E1 cysteine(s) closest to the contact region, the closest TFMD-tagged residues in CYP3A4, and potentially charge-pairing residues located in the contact zone identified by HEX docking. The passive residues were automatically selected within a 12 Å zone around the active residues. The remaining docking parameters were left at their default settings.

## 3. Results

### 3.1 Developing a high-throughput assay of 7-BQ debenzylation

7-BQ is widely used in the CYP3A activity assays. While the substrate itself exhibits a broad fluorescence band with emission and excitation maxima at 430 nm and 330 nm, respectively, its O-debenzylation shifts these maxima to 510 nm and 402 nm, respectively. The assay for 7-BQ O-debenzylation based on a time-dependent increase in fluorescence at 510 nm measured in a fluorimeter with a traditional optical cell and excitation at 400-405 nm is robust and sensitive [53, 54]. However, its high-throughput implementation for acquiring substrate-saturation profiles with a fluorescence plate reader encounters substantial challenges.

In our experience, all commercially available 7-BQ preparations are significantly (up to 0.5%) contaminated with the debenzylation product (7-HQ). As a result, simple measurements of fluorescence at 510 nm in a series of samples incubated with increasing concentrations of unpurified 7-BQ are prone to significant errors due to the fluorescence of the 7-HQ contaminant added along with the substrate. To address this issue, we developed a simple procedure for 7-BQ purification using silica solid-phase extraction columns, as described in Materials and Methods.

Although single-point fluorescence assays using re-purified 7-BQ with measuring emission at 510 nm with excitation at 405 nm may yield satisfactory results, the best accuracy and reproducibility are achieved when recording spectra with synchronous scanning of excitation and emission in 290 – 460 nm wavelength range with 110 nm wavelength difference and approximating them with a combination of prototypical spectra of fluorescence of 7-HQ, 7-BQ and NADPH. The detailed description of our setup is provided in the Materials and Methods section.

### 3.2. 7-BQ metabolism by recombinant P450 enzymes

The parameters of 7-BQ metabolism by individual P450 species obtained in our studies with Supersomes® containing recombinant P450 enzymes are shown in Table 1. As seen from these data, the highest efficiency in 7-BQ metabolism is exhibited by CYP3A4 and CYP3A5. Although quite a few other P450 species can metabolize the substrate, their affinity to 7-BQ and the maximal rate of its turnover are considerably lower than those exhibited by CYP3A enzymes.

**Table 1.** Parameters of 7-BQ O-demethylation by recombinant P450 enzymes in Supersomes.

| Cytochrome P450 species | $S_{50}$ or $K_M$, $\mu\text{M}$ | $N$ | $V_{\max}$ | $CL_{\text{int}}$, $\text{min}^{-1} \mu\text{M}^{-1}$ |
| --- | --- | --- | --- | --- |
| CYP3A4 | 43.9 ± 6.9 | 1.85 ± 0.24 | 108 ± 19 | 2.5 |
| CYP3A5 | 102 ± 11 | 1.28 ± 0.15 | 93 ± 39 | 0.91 |
| CYP1A2 | 125 ± 25 | | 30 ± 12 | 0.24 |
| CYP2B6 | 150 ± 29 | | 27 ± 13 | 0.18 |
| CYP2C19 | 45.9 ± 2.8 | | 6.9 ± 0.3 | 0.15 |
| CYP2D6 | 59.9 ± 1.9 | | 5.8 ± 0.8 | 0.10 |
| CYP2E1 | 197 ± 55 | | 8.1 ± 3.7 | 0.04 |
| CYP2C9 | 495 ± 193 | | 9.5 ± 2.8 | 0.02 |
| CYP2C8 | 441 ± 129 | | 8.1 ± 2.4 | 0.02 |

\* The values provided in the table represent the averages of the results of 4–7 individual experiments. The ± values correspond to the confidence intervals calculated for $p = 0.05$ . The $V_{\max}$ values are expressed as moles of 7-HQ formed per minute per mole of cytochrome P450 ( $\text{min}^{-1}$ ).

As illustrated by substrate-saturation profiles (SSP) of CYP3A4 and CYP3A5 with 7-BQ shown in Figure 1, both enzymes display noticeable homotropic cooperativity with this substrate. While the turnover rates of the two enzymes with 7-BQ are comparable, CYP3A4 exhibits higher affinity and greater cooperativity with the substrate than CYP3A5.

**Figure 1.**
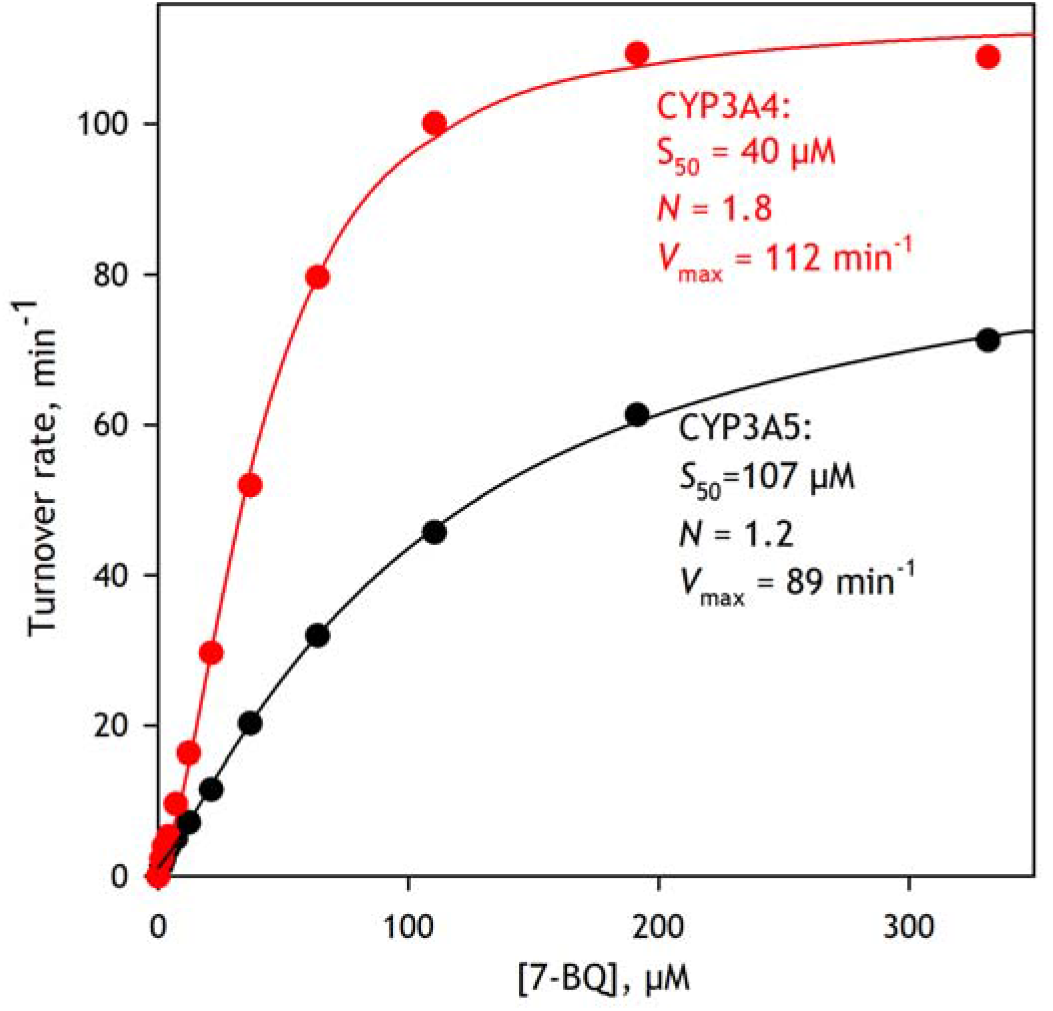
Substrate saturation profiles of 7-BQ debenzylation by CYP3A4 (red) and CYP3A5 (black) enzymes in Supersomes. The datasets represent averages of 5 experiments.

### 3.3. Metabolism of 7-BQ by HLM and the impact of chronic alcohol exposure of liver donors

A set of SSPs obtained with a series of 23 proteomically characterized HLM samples from donors with varying degrees of alcohol exposure, and the results of applying principal component analysis (PCA) to this dataset are shown in Figure 2. As illustrated in Figure 2a, the analyzed samples exhibit a wide variation in both SSP amplitudes and shapes.

**Figure 2.**
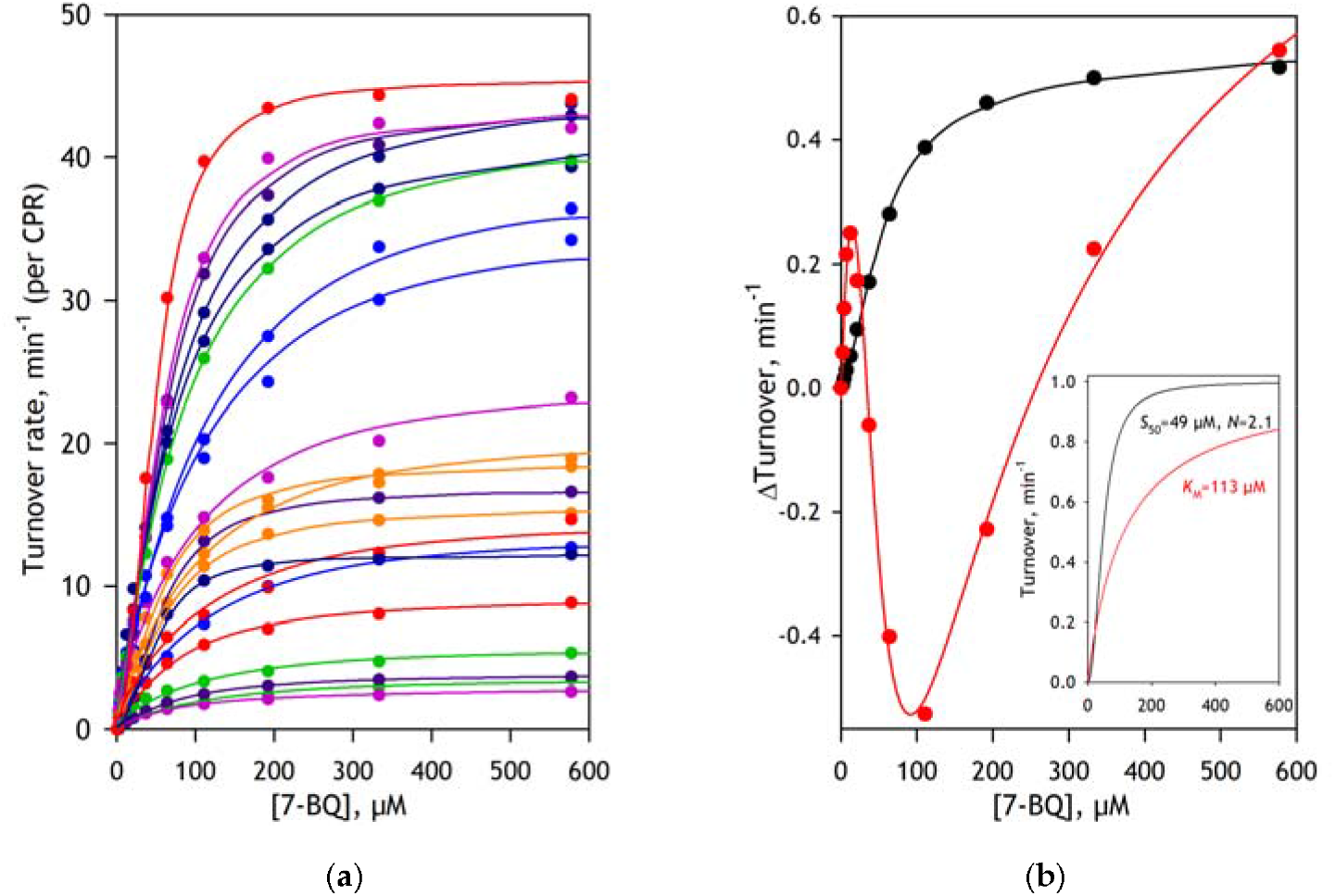
Substrate saturation profiles of O-debenzylation of 7-BQ and their analysis by PCA. Panel **a** shows a dataset obtained with a set of 23 individual HLM preparations. Each SSP shown in panel (***a)*** represents an average of the results of 2 - 4 individual experiments. The solid lines represent approximations of the SSPs as linear combinations of a Michaelis-Menten and a Hill equation, with globally optimized parameters. The results of applying PCA to this dataset are shown in panel (***b)***. The vectors of the first and second principal components are shown in the main panel in black and red, respectively. The solid lines in the main panel show their approximations as linear combinations of a Michaelis-Menten and a Hill equation, with globally optimized parameters shown in the insert.

The first two principal components resulting from PCA of this dataset (Figure 2b) account for over 99.5% of the observed differences. While the first component mainly reflects variation in SSP amplitudes, the second accounts for differences in the contributions of the higher- and lower-affinity components to the profiles. Inset in Figure 2b shows a pair of Hill- and Michaelis-Menten-equation components resolved by the PCA-based global fitting. Their linear combinations approximate all SSPs in this dataset with the square correlation coefficient (*R*^2^) not lower than 0.98 and, in most cases, higher than 0.995. As seen from this plot, the parameters of the higher-affinity component are in good agreement with those exhibited by CYP3A4, whereas the parameters of the low-affinity component are consistent with those of the two other most efficient 7-BQ metabolizers, CYP3A5 and CYP1A2.

We used the results of this global analysis to examine the impact of alcohol exposure on the parameters of 7-BQ metabolism. In this examination, we rely on the provisional index of alcohol exposure (PIAE) that we introduced in our recent study on a large collection of human liver samples [11]. This index grades alcohol exposure on a scale from 0 to 5 based on the relative abundances of four marker proteins (HSPA5, PDIA3, CES2, and P4HB) in the donor’s liver.

Fig. 3a shows the plots of the maximal rate of 7-BQ turnover (*V*_max total_) and the relative abundance of CYP2E1 (*RA*CYP2E1) versus PIAE. It demonstrates a strong correlation between the two parameters and the alcohol exposure. After removing the two extreme points on each side of the graph (the points with the two highest and two lowest PIAE values), the correlation between the rate of 7-BQ turnover and the level of alcohol consumption becomes well-pronounced. It is characterized by a correlation coefficient (*R*) of 0.58 and a T-test *p*-value of 0.01.

**Figure 3.**
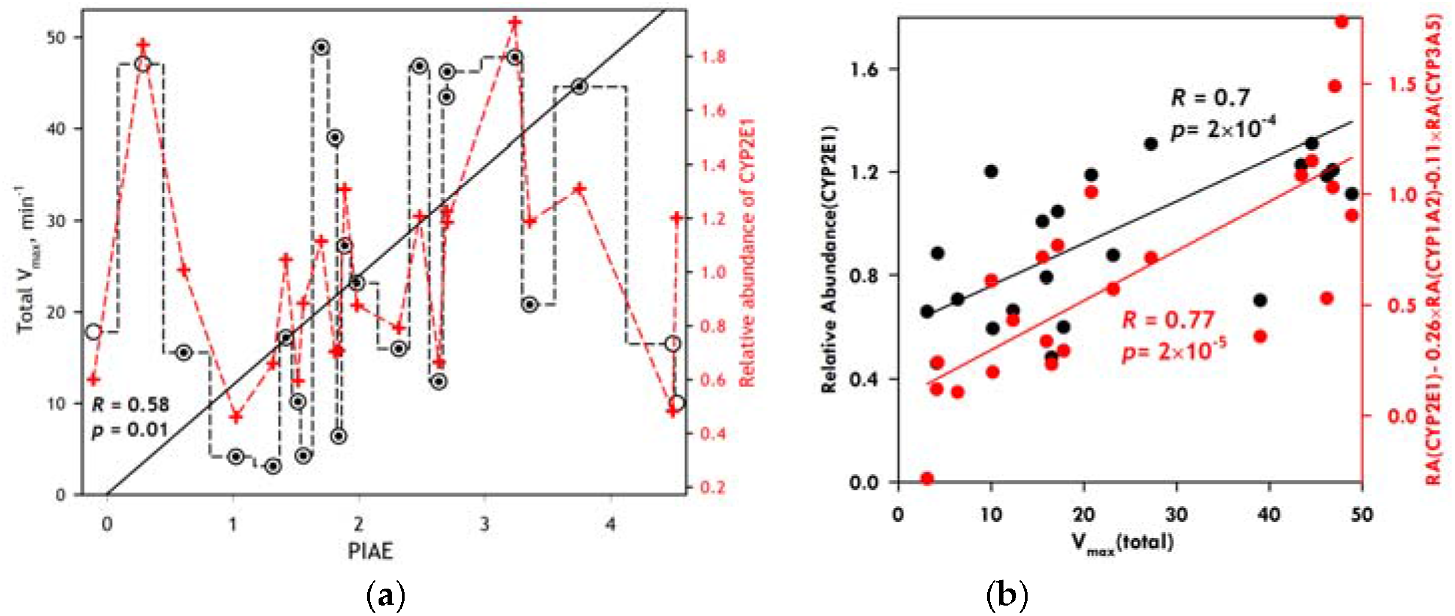
Correlation of the total maximal rate of BQ metabolism with the degree of alcohol exposure and relative abundance of CYP2E1. Panel (***a)*** shows the dependence of V_max total_ (open circles) and the relative abundance of CYP2E1 (red crosses) on PIAE. The solid line shows the linear approximation of 19 data points of the data set (dotted circles). Panel (***b***) shows the plots of the relative abundance of CYP2E1 (black) and the combination of the relative abundances of CYP2E1, CYP1A2, and CYP3A5 (red) versus V_max total_.

A comparison of the profiles of *V*_max total_ and *RA*_CYP2E1_ versus PIAE reveals an evident correlation between CYP2E1 abundance and the rate of BQ metabolism, characterized by *R* = 0.70 and a *p*-value of 2·10^-4^. This cannot be explained by the direct involvement of CYP2E1 in 7-BQ metabolism, since the turnover rate exhibited by CYP2E1 is many times lower than that of CYP3A4. At the same time, the concentrations of these two enzymes in HLM are comparable. It instead reveals an activating effect of CYP2E1 on CYP3A enzymes, in good agreement with our earlier observations [32].

To further investigate the correlations between the rate of 7-BQ metabolism and the composition of the P450 ensemble, we assessed the dependence of *V*_max total_ on PIAE by all possible combinations of two and three profiles of the relative abundance of 15 major P450 species, complemented with those of CPR and cytochrome *b*_5_. As illustrated in Fig. 3b, the best combination of three RA profiles comprises CYP2E1, CYP1A2, and CYP3A5, with the latter two providing negative contributions. This combination increases the correlation coefficient to 0.77 and yields a *p*-value as low as 2·10^-5^. The decrease in the rate of 7-BQ turnover at increased abundances of CYP3A5 and CYP1A2 is easily understandable, as these enzymes, while metabolizing 7-BQ at a lower rate, compete with CYP3A4 for interactions with CPR and for involvement in 7-BQ metabolism.

### 3.4. Effect of alcohol exposure on the metabolism of ivermectin

A bulky molecule of antiparasitic drug ivermectin (MW=875) is metabolized almost exclusively by CYP3A4 and CYP3A5 through 3”-O-demethylation and hydroxylation [55, 56, 57]. We used this substrate as an additional probe for the effect of alcohol exposure on CYP3A enzymes. Our high-throughput fluorimetric assay [39] enabled us to detect the formaldehyde produced during ivermectin demethylation.

Substrate-saturation profiles of ivermectin demethylation by CYP3A4 and CYP3A5 in Supersomes are shown in Fig. 4. As seen from this graph, the activity of CYP3A4 in O-demethylation of ivermectin is higher than that of CYP3A5, in agreement with earlier reports [55, 56]. However, the latter has a higher affinity to the substrate. In contrast to CYP3A4, which exhibits Michaelis-Menten kinetics, CYP3A5 shows a pronounced negative cooperativity.

**Figure 4.**
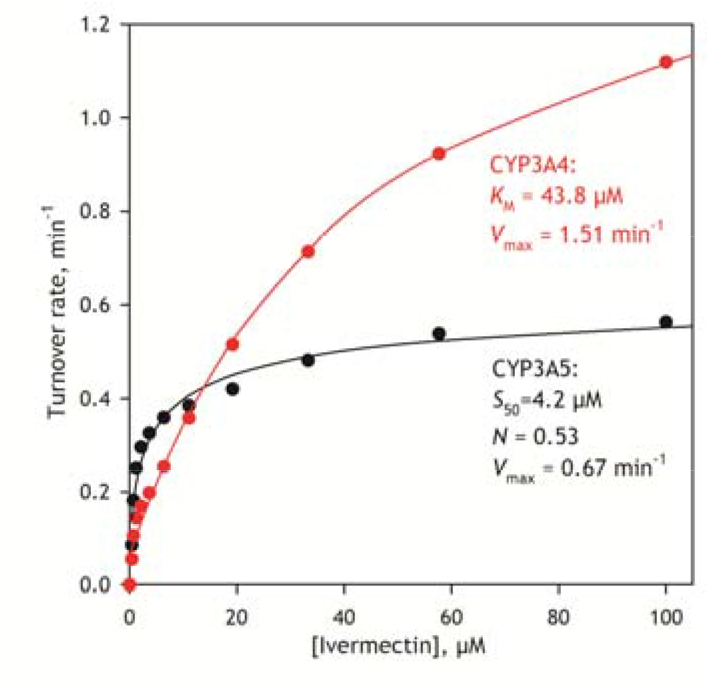
Substrate saturation profiles of ivermectin demethylation by CYP3A4 and CYP3A5 enzymes in Supersomes. The datasets represent averages across three experiments.

A set of SSPs obtained with our collection of HLM samples is shown in Figure 5A. As in the case of BQ, we observed considerable variation in the amplitude and shape of the profiles. PCA-assisted global fitting approximated the profiles as linear combinations of Michaelis-Menten dependence and a Hill equation (Figure 5B).

**Figure 5.**
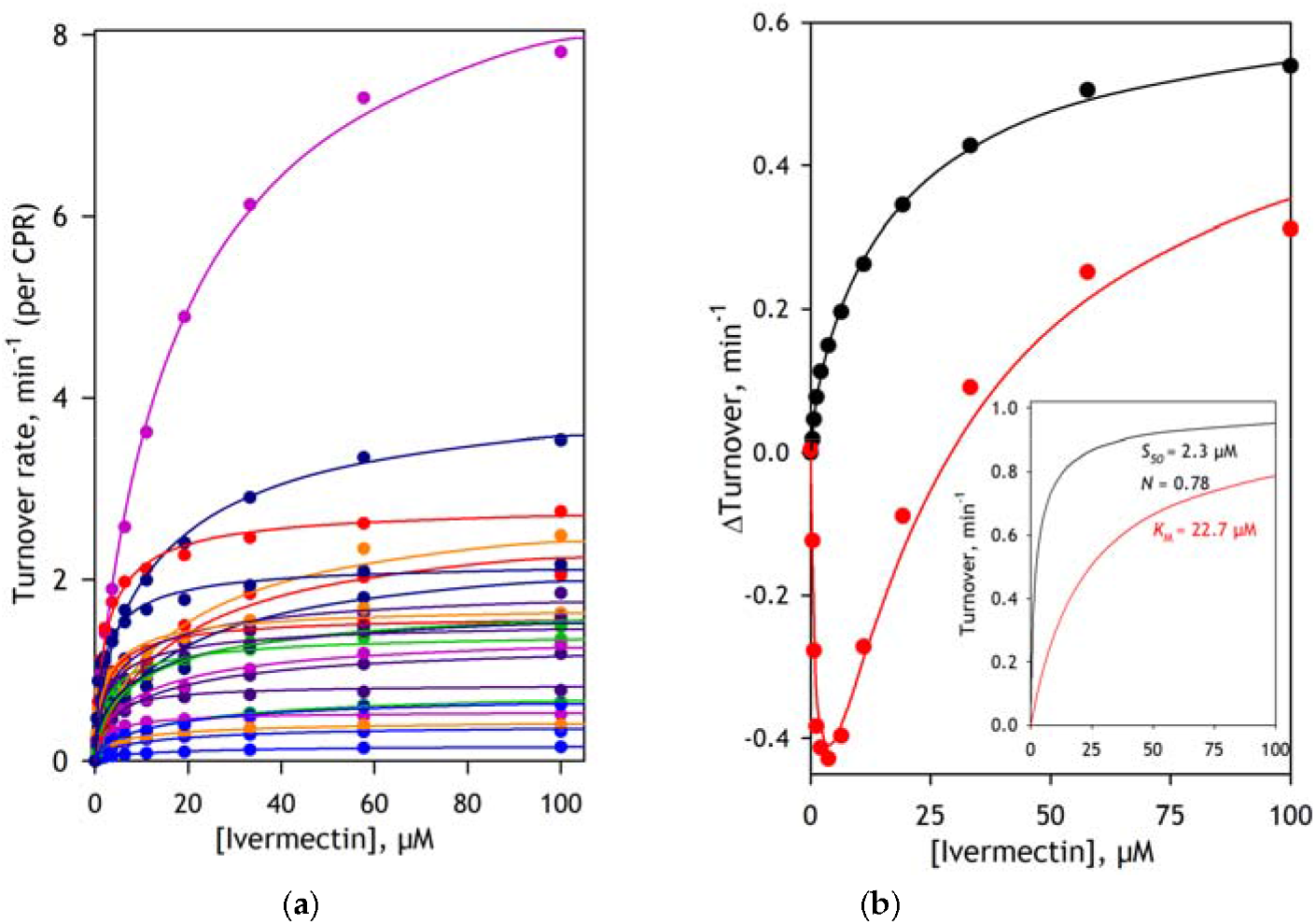
Substrate saturation profiles of O-demethylation of ivermectin and their analysis by PCA. Panel (***a***) shows a dataset obtained with a set of 23 individual HLM preparations. Each SSP shown in panels (***a***) represents an average of the results of 2 - 4 individual experiments. The solid lines represent approximations of the SSPs as linear combinations of a Michaelis-Menten and a Hill equation, with globally optimized parameters. The results of applying PCA to this dataset are shown in panel b. The vectors of the first and second principal components are shown in the main panel in black and red, respectively. The solid lines in the main panel show their approximations as linear combinations of a Michaelis-Menten and a Hill equation, with globally optimized parameters shown in the insert.

The dependence of *V*_max total_ on PIAE, shown in Fig. 6a, demonstrates a prominent increase in the turnover rate with increased alcohol exposure. Similar to the case of BQ, the correlation between *V*_max_total_ and PIAE becomes evident after removing two outliers from both the low and high ends of the graph. The dataset reduced in this way exhibits a correlation coefficient of 0.7 and a T-test *p*-value of 2·10^-3^.

**Figure 6.**
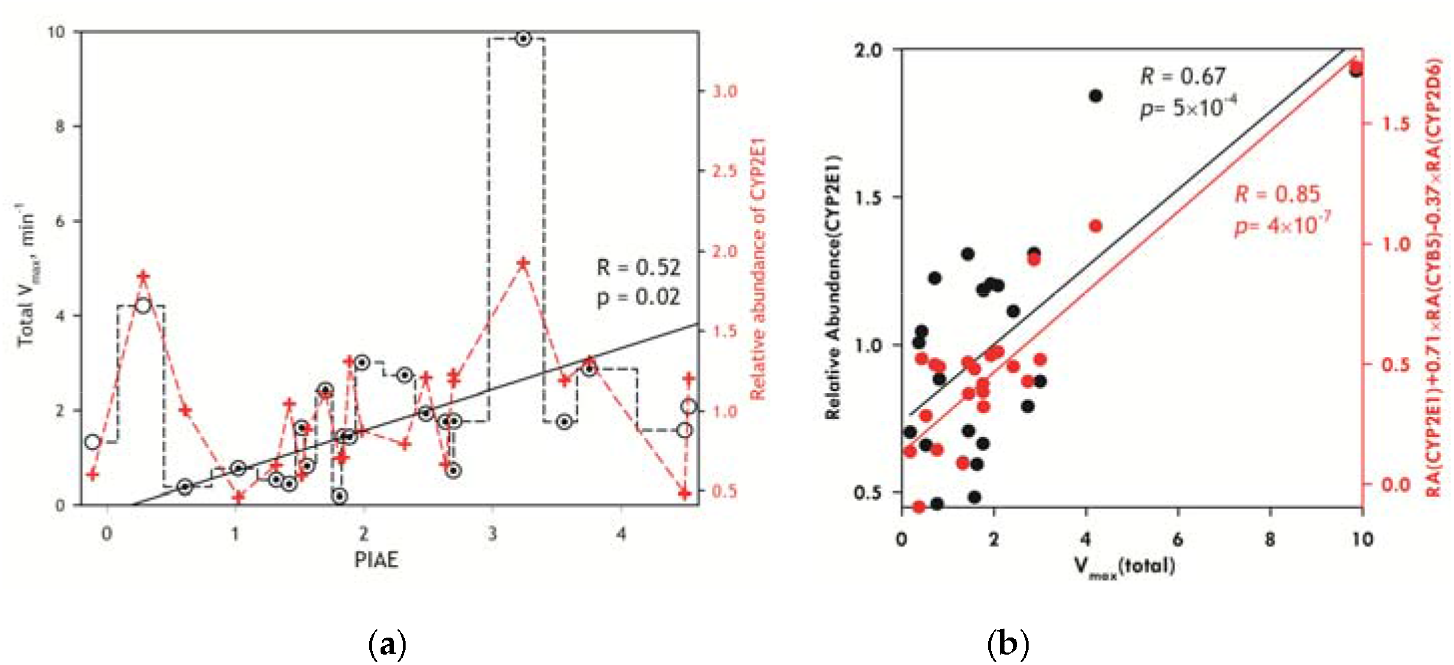
Correlation of the total maximal rate of ivermectin metabolism with the degree of alcohol exposure and relative abundance of CYP2E1. Panel (***a***) shows the dependence of *V*_max total_ (open circles) and the relative abundance of CYP2E1 (red crosses) on PIAE. The solid line shows the linear approximation of 19 data points of the data set (dotted circles). Panel (***b***) shows the plots of the relative abundance of CYP2E1 (black) and the combination of the relative abundances of CYP2E1, cytochrome b5, and CYP2D6 (red) versus *V*_max total_.

Similar to 7-BQ, the value of *V*_max total_ is correlated with CYP2E1 abundance across the full range of PIAE values in our dataset (Fig. 6B). This correlation exhibits *R* = 0.67 and a p-value of 5·10^-4^. No correlations between *V*_max total_ and the abundances of CYP3A4 or CYP3A5 were detected. However, the best combination of three profiles of relative abundance also reflects the positive contribution of cytochrome *b*_5_ (CYB5) and apparent inhibition (negative contribution) by CYP2D6. The combination of relative abundances of CYP2E1, cytochrome *b*_5_, and CYP2D6 correlates with *V*_max_total_ with a correlation coefficient of 0.85 and a *p*-value of 2·10^-7^ (Fig. 6b). The positive correlation between *V*_max total_ and the cytochrome *b*_5_ content is consistent with the well-known stimulating effect of cytochrome *b*_5_ on the activity of CYP3A enzymes [58, 59, 60]. In contrast, the reason for the apparent inhibitory effect of CYP2D6 remains unclear.

### 3.5. Probing the interactions of CYP2E1 with CYP3A4 with chemical crosslinking mass spectrometry (CXMS)

The activating effect of alcohol-inducible CYP2E1 on the activity of CYP3A4 described above was also evidenced in our earlier experiments with the incorporation of purified CYP2E1 into pooled HLM preparations [32]. These indications of the functional effects of CYP2E1-CYP3A4 interactions prompted us to probe the formation of CYP2E1-CYP3A4 complexes in microsomal membranes using chemical crosslinking mass spectrometry.

In these experiments, we combined the tag-transfer mass spectrometry approach with the molecular fishing strategy. As a cleavable photo-activated crosslinking agent, we used trifluoromethyl phenyl diazirine methanethiosulfonate (TFMD-MTS, 3-{4-[(methanesulfonylsulfanyl)methyl]phenyl}-3-(trifluoromethyl)-3H-diazirine) [45]. As a crosslinker-activated molecular bait, we employed purified His-tagged CYP2E1 protein modified with TFMD-MTS in the molecular ratio 1:3. The protein was incorporated into the membrane of insect cell microsomes (Supersomes®) containing recombinant CYP3A4, CPR, and cytochrome *b*_5_. After exposure to light, the membrane was solubilized, the bait was isolated by Ni-affinity chromatography, and the sample was subjected to SDS-PAGE. The fragments of the gel corresponding to molecular masses of 45 - 65 kDa (zone A), 65 – 150 kDa (zone B), and 150 – 300 kDa (zone C) were then analyzed by LC-MS/MS after trypsin digestion and cleaving S-S bonds of TFMD-MTS by alkylation (see Materials and Methods).

This analysis identified several TFMD-tagged peptide fragments of CYP3A4, suggesting a high degree of crosslinking between CYP2E1 and CYP3A4. The positions of the apparent sites of CYP2E1-CYP3A4 contacts revealed by these results are summarized in Table 2 and illustrated in Figure 7. As shown in this figure, the positions of the tagged residues in the CYP3A4 molecule suggest at least two distinct modes of interaction. Heavy tagging in the region of C and D α-helices at the proximal side of the protein is barely compatible within the same interaction mode with the contacts in the regions of the meander loop, beta-bulge, and B/B’ loop located at the distal side of the protein. These two distant contact regions may reveal two distinct types of CYP2E1-CYP3A4 interactions in their heteromers.

**Table 2.**
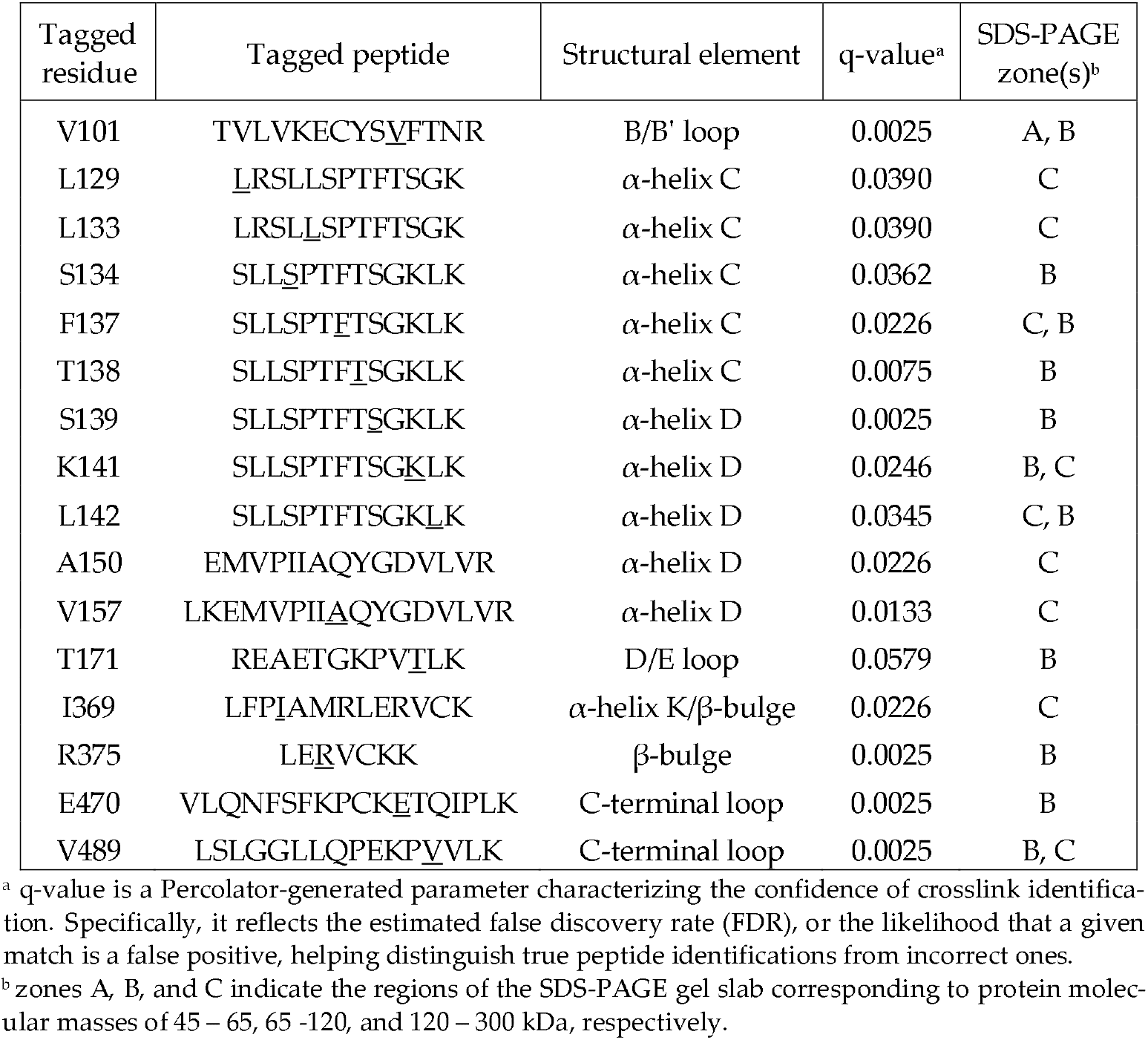
TFMD-tagged residues in CYP3A4 identified by tag-transfer CX-MS.

**Figure 7.**
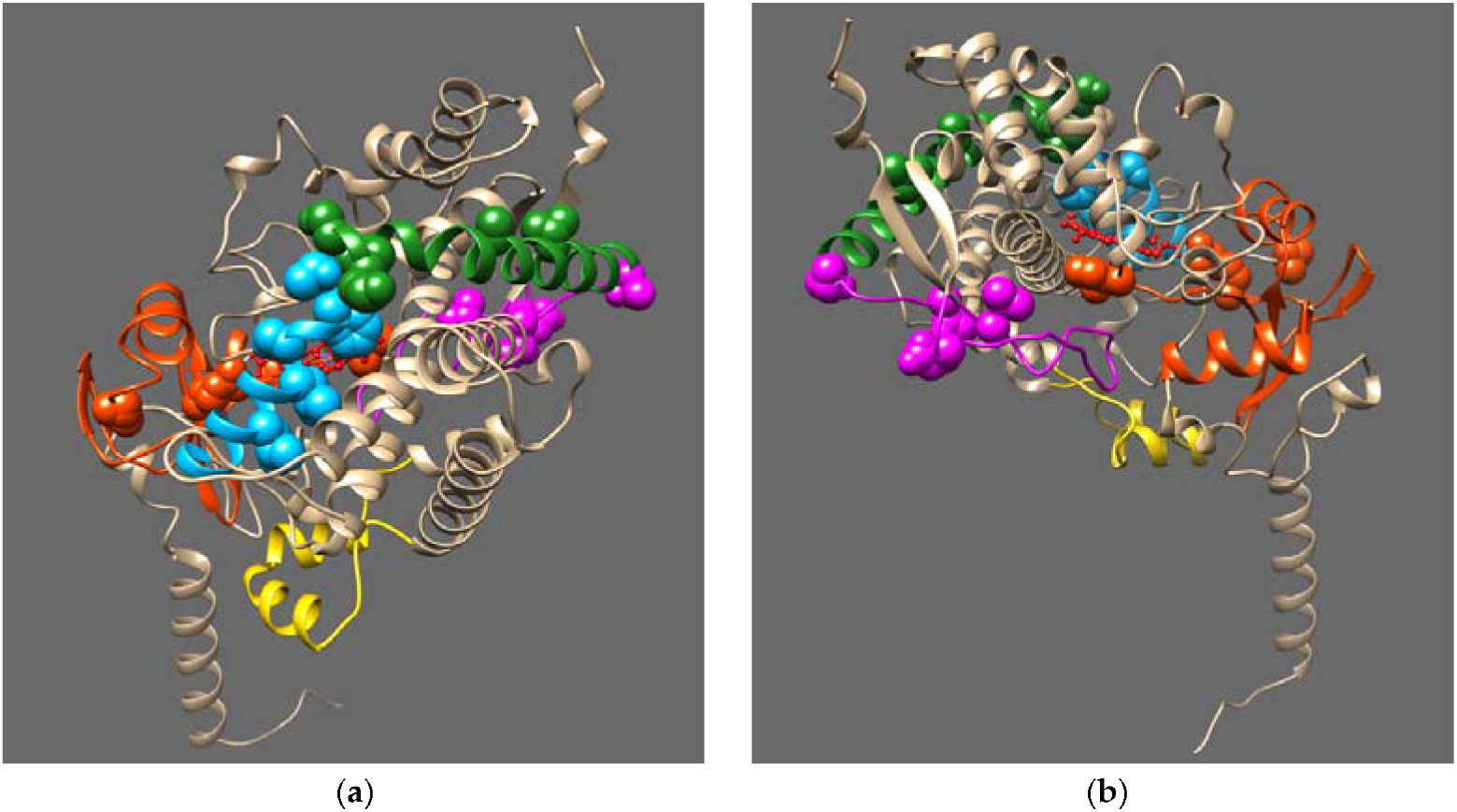
Location of TFMD-tagged residues in the CYP3A4 molecule. Panels (***a***) and (***b***) show the views of the molecule from the G-helix and A-helix sides, respectively. The tagged residues are shown in spherical atom models. Yellow, orange, blue, green, and magenta colors indicate the F/G-loop, beta-bulge/helix A/helix K region, helices C and D, and the C-terminal loop, respectively. The heme is shown as a red ball-and-stick model.

We attempted to model the architecture of CYP2E1-CYP3A4 complexes based on the locations of contact regions inferred from the positions of TFMD-tagged residues in CYP3A4. For this modeling, we assumed that, for successful crosslinking, the target residues in CYP3A4 must be within 20 Å of certain CYP2E1 cysteine residues accessible for modification with TFMD-MTS. In our protein docking trials, we used AlphaFold 2-generated models of full-length CYP2E1 and CYP3A4. Our docking procedure combined manual selection of a starting pose with subsequent restrained docking using HEX 8.0.0 [51, 52] followed by restrained docking using the HADDOCK 2.4 web server [49, 50]. The docking strategy is described in Section 2.9 of the Materials and Methods. The best docking poses produced at the final HADDOCK step were analyzed to select those that ensured proper orientation of the protein molecules in the membrane and proximity of at least one surface-exposed CYP2E1 cysteine (C261, C268, C452, C480, or C488) to the maximal possible number of TFMD-tagged residues in CYP3A4.

In our docking attempts, we probed eight different starting orientations for CYP2E1 and CYP3A4, including distal-to-distal, distal-to-proximal, proximal-to-distal, proximal-to-proximal, and four variants of side-to-proximal orientations. Our analysis indicates that the only cysteines of CYP2E1 that can be located in the proximity of the tagged regions of CYP3A4 for crosslinking with most of the tagged residues are the cysteines C261 and C268, as shown in Figure 8.

**Figure 8.**
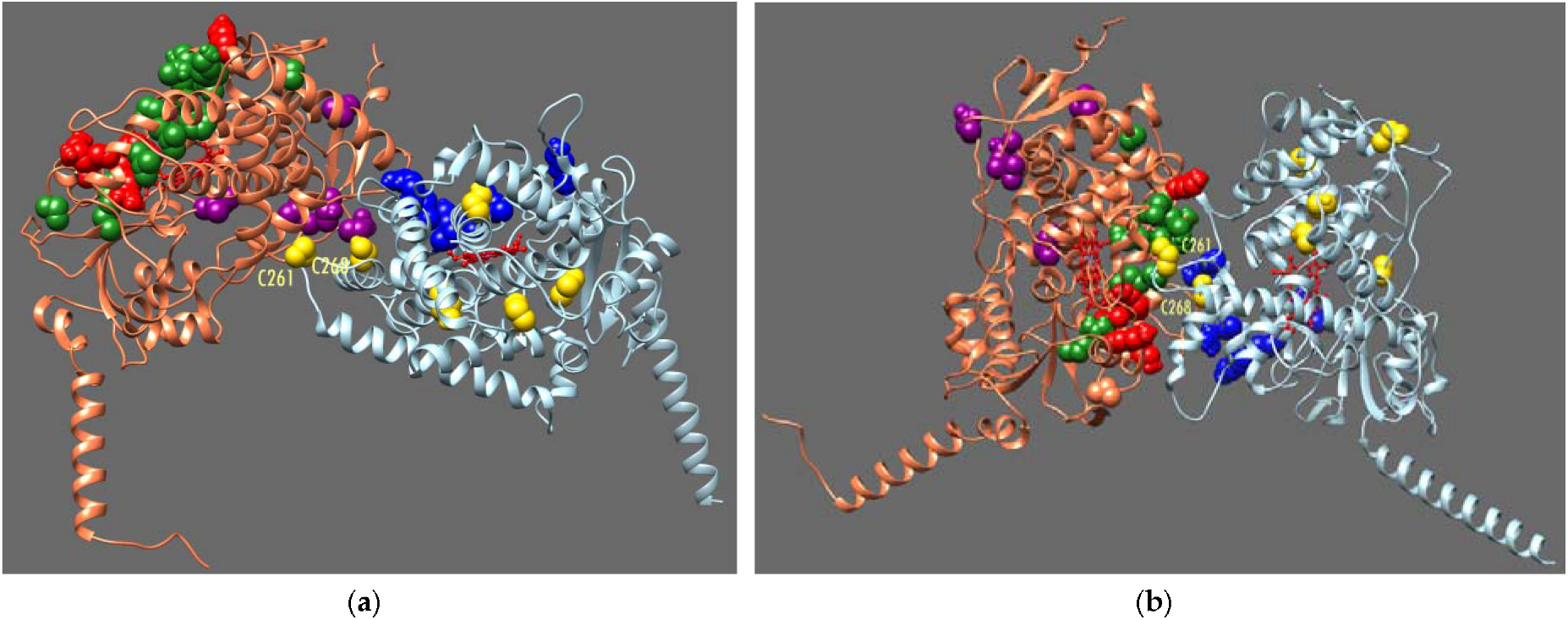
The two best models of CYP2E1-CYP3A4 complexes. Panel a shows Model 1, where CYP2E1 binds to the distal side of CYP3A4. Model 2, where the binding of CYP2E1 occurs at the proximal side of CYP3A4, is shown on panel b. TFMD-tagged residues in CYP3A4, located at the proximal and distal sides of the molecule, are shown as green and purple spherical atom models, respectively. CYP2E1 cysteine residues are shown in yellow spherical atoms. Red and blue spherical atoms indicate the residues presumably involved in interactions with CPR in CYP3A4 and CYP2E1, respectively. The heme groups are shown as red ball-and-stick models.

The two best docking positions are illustrated in Figure 8. They show distal (Model 1, Fig. 8a) and proximal (Model 2, Fig. 8b) CYP3A4 sides contacting the G-helix side of CYP2E1. The potential crosslinking pairs of residues for these models are summarized in Table 3. In Figure 8, the TFMD-tagged residues at the proximal and distal sides of CYP3A4 are shown in green and magenta spherical atom models, respectively.

**Table 3.** Parameters of the two best models of CYP2E1-CYP3A4 complexes.

| CYP3A4 residue | Model 1 (binding at the distal side of CYP3A4) |  | Model 2 (binding at the proximal side of CYP3A4) |  |
| --- | --- | --- | --- | --- |
|  | CYP2E1 residue | Distance, Å | CYP2E1 residue | Distance, Å |
| L129 |  |  | C268 | 13.4 |
| L133 |  |  | C261 | 11.4 |
| S134 |  |  | C268 | 8.2 |
| F137 |  |  | C261 | 5.25 |
| T138 |  |  | C268 | 8.5 |
| S139 |  |  | C261 | 4.3 |
| K141 |  |  | C261 | 10.2 |
| L142 |  |  | C261 | 5.5 |
| A150 |  |  | C261 | 19.4 |
| V157 | C268 | 20 |  |  |
| T171 | C261 | 10.8 |  |  |
| I369 | C261 | 15.9 |  |  |
| R375 |  |  | C261 | 17.3 |
| E470 | C268 | 12 |  |  |
| V489 | C268 | 13.8 |  |  |
| Haddock score | -92.6 |  | -118.8 |  |
| CPR binding site in CYP3A4 | open |  | obstructed |  |
| CPR binding site in CYP2E1 | open |  | open |  |

In these figures, we also show the positions of charge-pairing residues presumed to be involved in the interactions between CYP3A4 and CYP2E1 and CPR, represented as red and blue spherical atom models, respectively. These residues, namely K91, K127, R130, K143, and R440 in CYP3A4 and K87, K123, R127, R134, and K434 in CYP2E1 were identified using the models of the CYP2D6-CPR complex [61, 62] and the available data on CYP2B4-CPR interactions [63] based on sequence alignment of CYP2E1 and CYP3A4 with CYP2D6 and CYP2B4. The list of interacting residues was further refined based on models of the CYP3A4 complex with the FMN domain of CPR described by Urban and co-authors [60] and available at Model Archive (https://modelarchive.org) under identifiers ma-h9omo and ma-bihdn. As seen in Figure 8, the CPR binding site in CYP3A4 in Model 1 is open to interactions, whereas in Model 2, the binding of CPR to CYP3A4 is obstructed.

## 4. Discussion

In this study, we used a combination of high-throughput kinetic assays and global proteomic analysis to explore the effect of alcohol exposure on the activity of CYP3A enzymes in human liver and its correlation with alcohol-induced changes in the composition of the P450 ensemble in HLM. Our studies with two CYP3A-specific substrates, 7-BQ and ivermectin, revealed a striking increase in the activities of CYP3A4 and CYP3A5 enzymes caused by chronic alcohol exposure. However, this activation of CYP3A enzymes in alcohol consumers is not associated with changes in the levels of CYP3A enzymes, which do not reveal any correlation with alcohol exposure. Instead, we found that the rates of CYP3A-dependent metabolism of 7-BQ and ivermectin correlate with CYP2E1 content. However, this alcohol-inducible enzyme is a very poor metabolizer of 7-BQ and does not metabolize ivermectin. These results suggest that the reported alcohol-induced increase in the metabolism of CYP3A drug substrates, such as diazepam [3, 17], carbamazepine [26], opiates [21, 22, 23], or tetracycline ics [16], is, at least in part, caused by the functional effects of CYP3A interactions with CYP2E1. This conclusion aligns well with our earlier studies, in which we observed a boost in 7-BQ metabolism after HLM enrichment with CYP2E1 by incorporating purified protein [32].

Lack of correlation between the abundance of CYP3A4 in HLM and its activity in the metabolism of our probe substrates, which is instead tightly correlated with the content of alcohol-inducible CYP2E1, confirms our notion of a lack of straight additivity in the functional properties of the members of the microsomal P450 ensemble [30, 35]. These results are in good agreement with the concept of functional integration of multiple P450 species via their protein-protein interactions and the formation of heteromeric P450 complexes in the microsomal membrane [30, 34, 64, 65]. According to this concept, a significant fraction of the cytochrome P450 molecules in HLM is found in inactive (“latent”) positions within heteromeric complexes of multiple P450 species. Availability of a particular cytochrome P450 for interaction with substrates and reductase and, thus, its involvement in catalytic activity is a complex function of the preferences of individual species for occupying the “active” and “latent” positions, the composition of the P450 pool, and the presence of selective substrates of individual P450 species.

Understanding these mechanisms requires in-depth knowledge of the structural basis of P450-P450 interactions. The only available inferences on the architecture of homo- and heterooligomers of P450 species were based on the analysis of crystallographic multimers of several P450 enzymes [66]. Our tag-transfer CXMS studies of the CYP2E1-CYP3A4 complex provide the first direct information on the architecture of P450 heteromeric complexes, shedding light on possible mechanisms underlying the functional effects of P450-P450 interactions.

Our results align well with our earlier inference on two different modes of protein-protein interactions in CYP3A4 oligomers [67]. Mapping the CYP3A4 residues located in the CYP2E1-CYP3A4 contact regions revealed two distinct modes of interaction between the proteins, differing in the accessibility of CYP3A4 to CPR binding. In Model 2, where CYP2E1 interacts with the proximal side of CYP3A4, the CPR binding site in CYP3A4 is obstructed, whereas CYP2E1 retains the ability to bind the electron donor in both models. Notably, probing the binding of CPR to CYP3A4-interacting CYP2E1 using the models of CYP3A4 complex with FMN-domain of CPR [60], we found that for both Model 1 and Model 2, the only possible mode of CYP2E1-CPR interactions is similar to Type 1 (doi: 10.5452/ma-h9omo) described by Urban and co-authors [60]. The interactions in the Type 2 mode (doi: 10.5452/ma-bihdn) are hindered by the CYP2E1-bound CYP3A4 molecule.

We may hypothesize that the two types of subunit interactions in CYP3A4 homooligomers [67] are analogous to those seen in CYP3A4-CYP2E1 complexes, and some part of CYP3A4 in its homooligomers is blocked from the interactions with CPR due to its involvement in the side-to-proximal complexes (similar to Model 2) between two CYP3A4 molecules. Hypothetical mechanism of activation of CYP3A4 by CYP2E1 may be based on preferential formation of the Type 1 CYP2E1-CYP3A4 complexes that would recruit additional CYP3A4 molecules into open positions. However, an in-depth understanding of this mechanism requires additional data on the architecture of both CYP3A4 homooligomers and its heterooligomers with CYP2E1.

Regardless of the mechanistic basis of the activating effect of alcohol-induced expression of CYP2E1 on CYP3A enzymes, it has a critical importance for practical pharmacology as it affects the pharmacokinetics of a plethora of drugs on the market, including those used for the treatment of alcohol use disorders (AUD) and alcohol withdrawal syndrome (AWS). A similar type of functional effects leading to ADU may occur in other P450 species, particularly CYP1A2 [34]. These potential ADU-related functional interactions in the P450 ensemble are currently under active investigation in our laboratories.

## Acknowledgements

The authors gratefully acknowledge Dr. Michael R. Hoopmann and Dr. Michael Riffle (Institute for Systems Biology, University of Washington, Seattle, WA) for their valuable technical insights into CX-MS data analysis performed using Kojak and ProXL.

## Author Contributions

Conceptualization, D.R.D.; methodology, D.R.D., B.P., D.K.S., and A.G.N.; software, D.R.D..; formal analysis, D.R.D., D.K.S.. and A.G.N.; investigation, D.R.D., K.P., N.D., K.A.G., and Y.G.; resources, D.R.D. and B.P.; writing—original draft preparation, D.R.D., K.P. and D.K.S..; writing—review and editing, D.R.D. and B.P..; supervision, D.R.D. and B.P..; funding acquisition, D.R.D. and B.P. All authors have read and agreed to the published version of the manuscript.

## Institutional Review Board Statement

Ethical review and approval were waived for this study because it was performed with IRB-exempt deidentified tissue specimens.

